# The case for using Mapped Exonic Non-Duplicate (MEND) read counts in RNA-Seq experiments: examples from pediatric cancer datasets

**DOI:** 10.1101/716829

**Authors:** Holly C. Beale, Jacquelyn M. Roger, Matthew A. Cattle, Liam T. McKay, Drew K. A. Thomson, Katrina Learned, A. Geoffrey Lyle, Ellen T. Kephart, Rob Currie, Du Linh Lam, Lauren Sanders, Jacob Pfeil, John Vivian, Isabel Bjork, Sofie R. Salama, David Haussler, Olena M. Vaske

## Abstract

**Background:** The accuracy of gene expression as measured by RNA sequencing (RNA-Seq) is dependent on the amount of sequencing performed. However, some types of reads are not informative for determining this accuracy. Unmapped and non-exonic reads do not contribute to gene expression quantification. Duplicate reads can be the product of high gene expression or technical errors.

**Findings:** We surveyed bulk RNA-Seq datasets from 2179 tumors in 48 cohorts to determine the fractions of uninformative reads. Total sequence depth was 0.2-668 million reads (median (med.) 61 million; interquartile range (IQR) 53 million). Unmapped reads constitute 1-77% of all reads (med. 3%; IQR 3%); duplicate reads constitute 3-100% of mapped reads (med. 27%; IQR 30%); and non-exonic reads constitute 4-97% of mapped, non-duplicate reads (med. 25%; IQR 21%). Informative reads--Mapped, Exonic, Non-duplicate (MEND) reads--constitute 0-79% of total reads (med. 50%; IQR 31%). Further, we find that MEND read counts have a 0.22 Pearson correlation to the number of genes expressed above 1 Transcript Per Million, while total reads have a correlation of −0.05.

**Conclusions:** Since the fraction of uninformative reads vary, we propose using only definitively informative reads, MEND reads, for the purposes of asserting the accuracy of gene expression measured in a bulk RNA-Seq experiment. We provide a Docker image containing 1) the existing required tools (RSeQC, sambamba and samblaster) and 2) a custom script. We recommend that all results, sensitivity studies and depth recommendations use MEND units.

## Background

Assessing the accuracy and reproducibility of gene expression results obtained from the analysis of RNA-Seq data has been a priority since the development of the assay. Seminal studies showed the relationship between the amount of sequence data generated during an experiment (depth of sequencing) and the reproducibility of the resulting gene expression measurements [1,2]. However, RNA-Seq data is not homogenous. Of the tens of millions of sequences (reads) in a typical RNA-Seq dataset (the data generated from one biological sample), some reads cannot be mapped back to the reference transcriptome. Others map to genome regions outside of exons or have been duplicated during the library construction process or sequencing. Nearly all methods for quantifying gene expression in bulk RNA-Seq data count reads that align to exons in a gene; thus, unmapped and non-exonic reads do not contribute to measurements and are uninformative regarding the accuracy of the experiment [3,4]. Therefore, considering the total number of reads as a proxy for RNA-Seq gene expression accuracy can result in inflated accuracy estimates.

Duplicate reads may be due to either highly abundant transcripts or technical error. The process of preparing RNA-Seq libraries involves PCR amplification. This step can generate duplicated identical or nearly identical reads. While the original read represents gene expression in the experimental system, the artifactual duplicate reads do not. However, duplicate reads are also generated by very highly expressed genes since each gene has a finite number of unique read sequences that can be generated from it [5].

Here we analyze 2179 bulk, paired end, polyA-selected RNA-Seq datasets to characterize the read types present in the datasets and evaluate what fraction of commonly reported data is unequivocally relevant to the accuracy of gene expression measurements. We compare the correlation of total reads and MEND reads to the number of measured genes.

## Methods

### MEND read counting method

Quantification of Mapped, Exonic, Non-Duplicate (MEND) reads was previously described [6]. Briefly, input is a genome-aligned bam file containing RNA-Seq read data. Duplicates are marked with Samblaster v0.1.22 [7], and the RSeQC v2.7.10 [8] script read_distribution.py quantifies exonic read and tag counts, excluding QC fail and duplicate reads as well as secondary alignments. The script parseReadDist.R, which we wrote, estimates the number of MEND reads based on RSeQC output by summing the tag counts in CDS exons, 5’ UTR exons and 3’ UTR exons and multiplying by reads per tag. Since a pair of reads provides information about two nearby sequences in a single transcript, read counts are reported in pairs. For example, 20 reads means that there are 20 pairs of reads. The process for estimating MEND read counts is available as a stand-alone docker image [9] and can be executed on CodeOcean [10]. The source code is freely available on GitHub [11].

### Data description

Here we discuss 2179 publicly available, bulk RNA-Seq datasets we gathered for the RNA-Seq compendium [12] used for comparative single-patient analysis [6]. Accession numbers, clinical data and read counts for each dataset are in Table S1. Repositories and cohort information is aggregated in Tables S2 and S3.

Of the 2179 datasets, 2018 were from pediatric/adolescent/young adult cancer tumors, 66 were from adult cancer tumors, and 95 were from cancer tumors of individuals with unknown ages, where adults are defined as being over 30 years of age. Of the 1692 datasets with reported gender of the patient, 42% were female and 58% were male. Of the 602 datasets with reported race of the patient, 27 patients were Asian, 70 were Black/African American, 3 were Native Hawaiian or Other Pacific Islander, 494 were White and 7 were Other without further definition. None were American Indian or Alaskan Native. Of 861 datasets with reported results of the patient’s Hispanic or Latino identity, 128 were Hispanic or Latino. The source tumors represent a variety of hematologic and solid malignancies (Table 1).

**Table 1:** Diseases present in studied datasets

| Disease | n | percent |
| --- | --- | --- |
| Acute lymphoblastic leukemia | 680 | 31.2% |
| Acute myeloid leukemia | 221 | 10.1% |
| Medulloblastoma | 201 | 9.2% |
| Glioma | 193 | 8.9% |
| Osteosarcoma | 157 | 7.2% |
| Acute megakaryoblastic leukemia | 103 | 4.7% |
| Ependymoma | 98 | 4.5% |
| Ewing sarcoma | 70 | 3.2% |
| Rhabdoid tumor | 65 | 3.0% |
| Rhabdomyosarcoma | 53 | 2.4% |
| Lymphoma | 49 | 2.2% |
| Embryonal rhabdomyosarcoma | 42 | 1.9% |
| Alveolar rhabdomyosarcoma | 40 | 1.8% |
| Glioblastoma multiforme | 29 | 1.3% |
| Choroid plexus carcinoma | 25 | 1.1% |
| Synovial sarcoma | 22 | 1.0% |
| Other | 131 | 6.0% |

The datasets came from five repositories (Table S2). Each was assigned to a cohort based on 1) project accession (for EGA and SRA datasets), 2) disease sub-study for NCI Therapeutically Applicable Research to Generate Effective Treatments (TARGET), or 3) disease for datasets in the St Jude Cloud. Cohorts were assigned IDs in descending order of size. Cohort assignments were intended to approximate a typical sequencing project performed by one research group at one sequencing center. The cohorts range in size from 3 to 337 datasets (Figure 1A); the median number of datasets in a cohort is 24.5.

**Figure 1:**
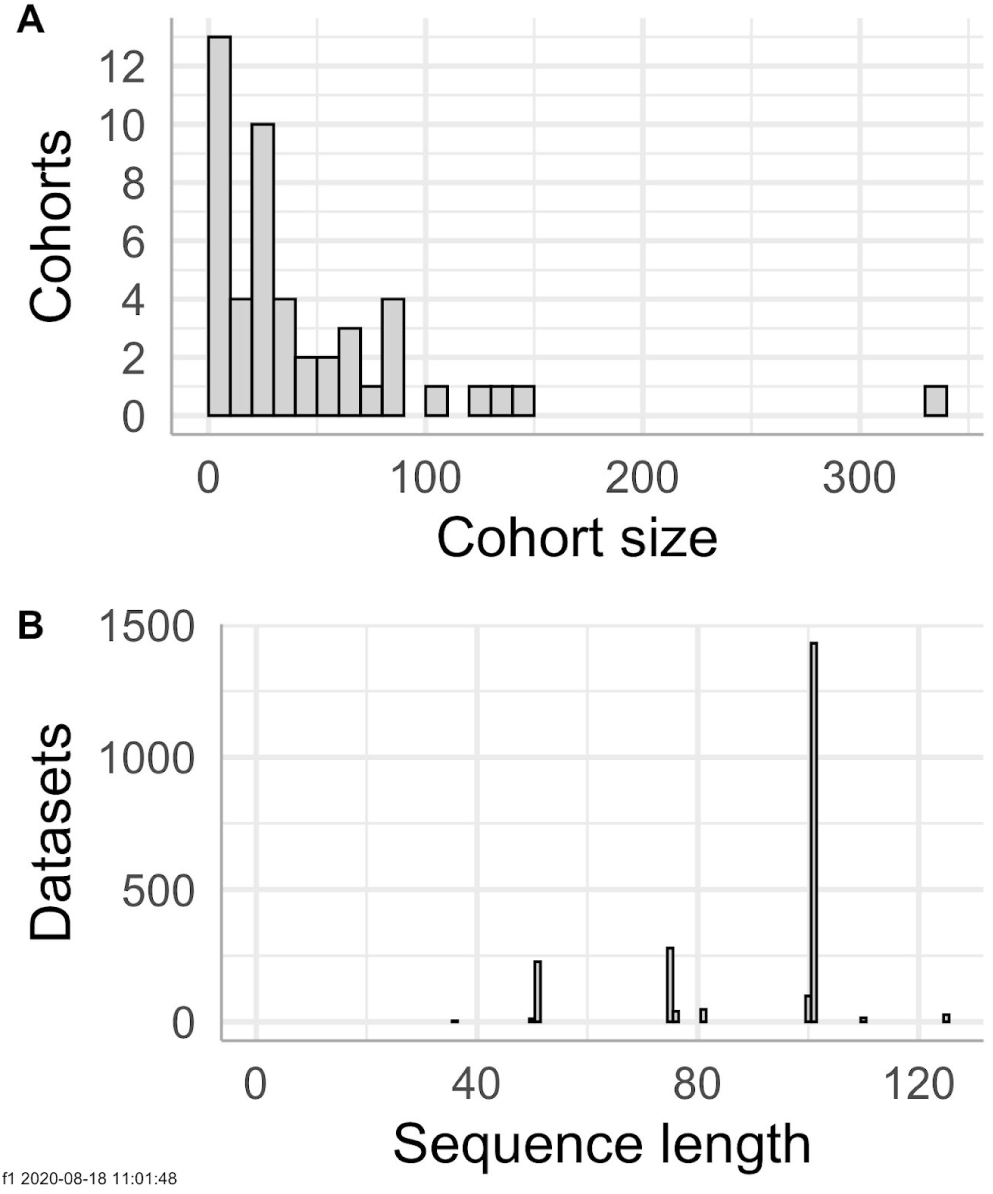
RNA-Seq datasets from 48 cohorts with a variety of read lengths were analyzed. A. Distribution of number of datasets per cohort. B. Distribution of length of paired end reads in this study.

All libraries were prepared with polyA selection. All data were generated via paired-end Illumina sequencing technology. The median sequence length is 101 bases (Figure 1B).

### Data analysis

RNA-Seq read data was aligned to the genome with the TOIL RNA-Seq pipeline previously described [13]. Briefly, adapters were removed with CutAdapt v1.9. Reads were then aligned with STAR v2.4.2a with indices based on GRCh38 and gencode v23. RSEM v1.2.25 was used to quantify gene expression. The source code of the pipeline is available [14]. MEND read counts were calculated with MEND qc release v1.1.1.

Read count and gene expression analysis was conducted with the R programming language, using the following packages: tidyverse, janitor, knitr, corr, cowplot, RColorBrewer, pander, kableExtra, and snakecase [15–24].

## Results

### Read types in RNA-Seq data

We interrogated the read types present in our RNA-Seq datasets in our gene expression quantification pipeline (Fig 1A). We obtained the number of total and mapped reads from the aligner log. We marked duplicates in the aligned BAM file, and counted them, along with exonic reads, using RSeQC. Duplicate reads are reported as a fraction of mapped reads, and exonic reads are reported as a fraction of non-duplicate reads.

Most RNA-Seq datasets contain a small percentage of unmapped reads (Fig 2B). In the data from 2179 datasets, 75% of datasets have fewer than 6% unmapped reads (Fig 3A). The distribution is left-skewed with a long right tail. The value of excluding these from read counts is apparent, as these reads do not correspond to any known expressed gene; in 77 datasets, more than 25% of reads are unmapped. Including those reads in any measure of the sensitivity of gene expression measurement would misguide the researcher.

**Figure 2:**
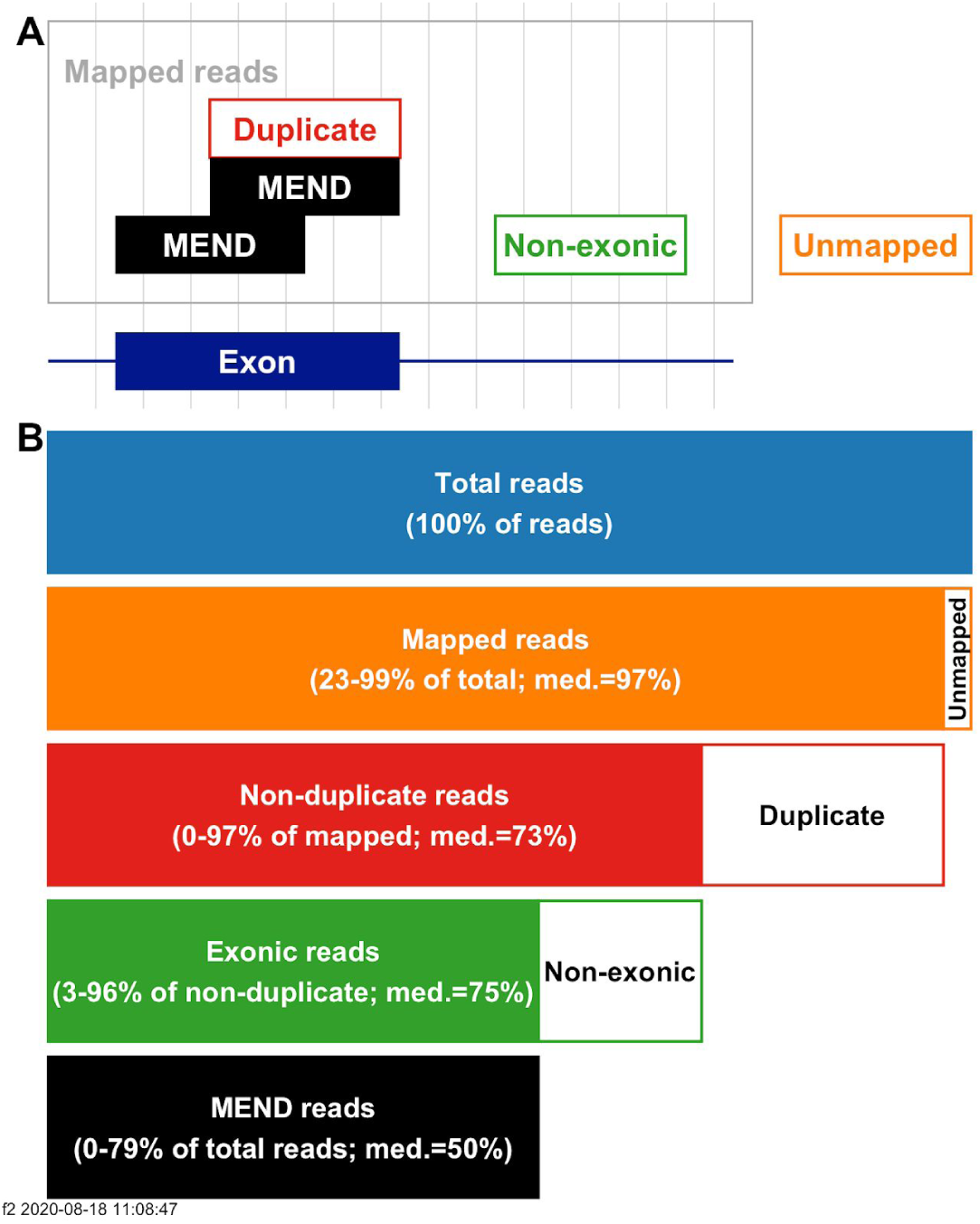
RNA-Seq datasets consist of 4 main types of sequencing reads. A. Simplified schematic illustrating read types. The X axis (blue) is a genomic sequence containing an exon. The other boxes each represent one sequencing read. Two of five reads are MEND reads. Other reads do not map to the genome (Unmapped; orange border), map to a non-exonic region of the genome (Non-exonic; green border), or are duplicates of other reads (Duplicate; red border). The MEND reads (black) fit none of these categories and are considered informative for determining the accuracy of gene expression quantification. B. Schematic illustrating read type quantification. Bars representing uninformative reads are white with a colored border. For each informative fraction, the range and median (med.) are reported.

**Figure 3:**
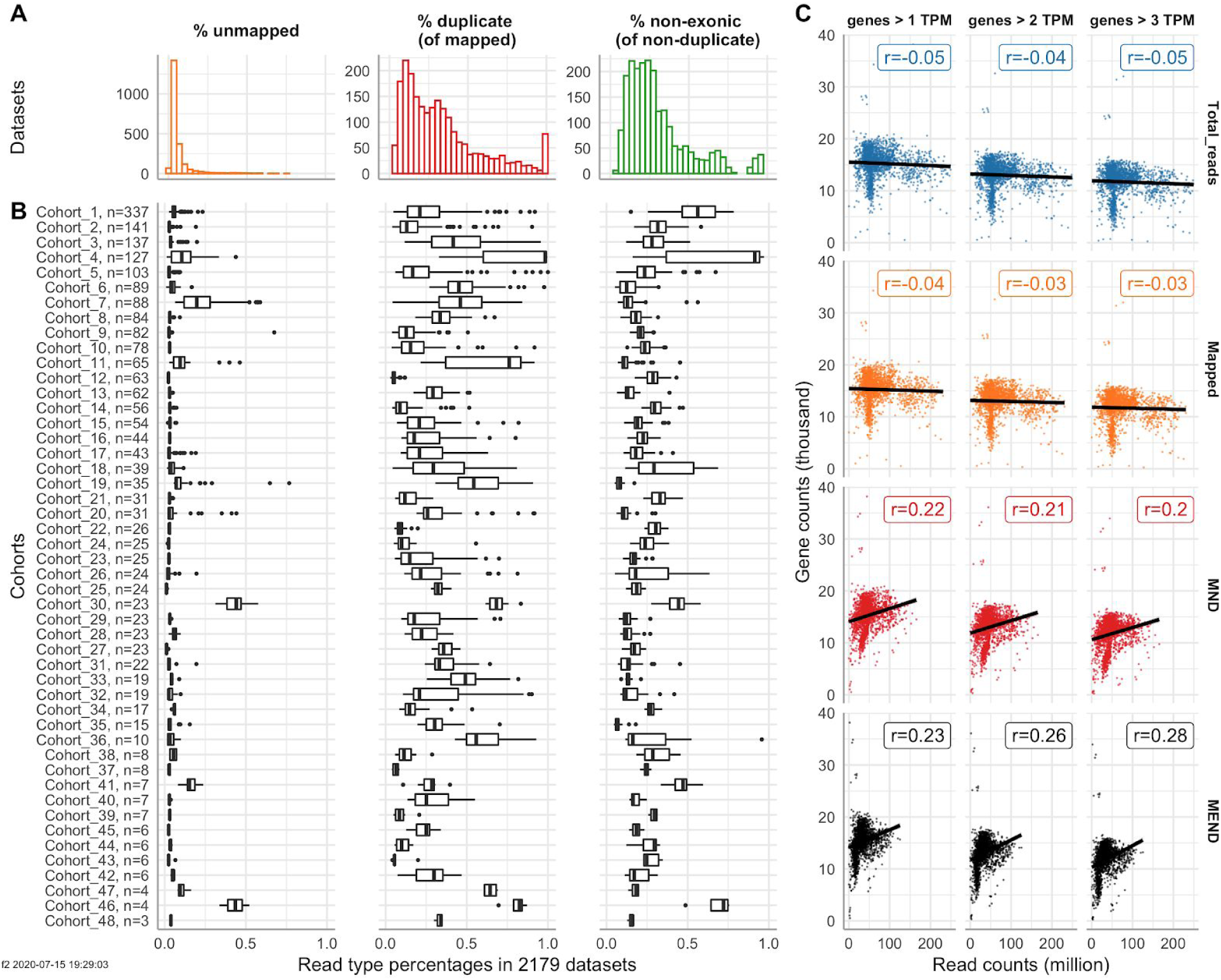
Read type fractions vary within and between cohorts, and MEND counts correlate best with measured gene counts. A. The percent distribution of different uninformative read types observed in 2179 datasets. B. The percentage of read types observed in cohorts, annotated with the number of datasets in the cohort. C: Relationships between the number of genes expressed in a dataset to the number of reads of different types. Only the 1996 datasets with more than 100 measured genes and fewer than 250 million total reads were included. Correlations to the number of genes expressed above 1, 2 and 3 TPM are shown.

The percentage of mapped reads that are duplicate reads (“percent duplicates”) is more varied. 426 datasets have more than 50% duplicates (Fig 3A). Some cohorts are characterized by high duplicate fractions (Fig 3B). For example, 72 of the 127 datasets in Cohort 4 have more than 98% duplicates. All 72 have a total sequencing depth above 170 million reads. However, even cohorts with generally low duplicate fractions can contain anomalous datasets; of the 41 cohorts with a median of less than 50% duplicates, 26 contain at least one dataset with more than 50% duplicates.

If duplicate reads were only a function of genes being especially deeply sequenced, we would expect sequencing depth to explain most of the variability in the fraction of duplicate reads. The total sequencing depth has a 0.58 Pearson correlation with the fraction of duplicate reads, explaining 34% of the variability (Supplemental Figure 1). The majority of the variability in the fraction of duplicate reads is independent of read depth. Consequently, the fraction of duplicate reads cannot be inferred from the total read depth.

Like percent of duplicates, the percent of non-exonic reads among all mapped, non-duplicate reads (“percent non-exonic”) has a broad distribution compared to other read type fractions, with an IQR of 21%. 330 datasets have a fraction of non-exonic reads above 50%.

### Correlation of read counts with the number of genes measured

If total or all mapped read counts were informative about the sensitivity of gene expression measurements, we would expect them to correlate with the number of expressed genes. When calculated using all 2179 datasets, total and mapped reads were inversely correlated (Pearson r=-0.4 for both) with the number of genes with expression above 1 Transcripts Per Million (TPM) (Supplemental Figure 2). We recalculated the correlations after excluding 1) the 78 datasets with fewer than 100 measured genes (all of which had more than 170 million total reads) and 2) the 105 datasets sequenced to more than 250 million total reads (such deep sequencing is usually intended for detecting rare events rather than measuring gene-level expression). In the resulting data from 1996 datasets, total and mapped reads are not correlated with the number of genes with expression above 1 TPM (Pearson r = −0.05 and -.0.04, respectively; Figure 2C). MND and MEND read counts are correlated at Pearson r = 0.22 and 0.23, respectively. When genes with higher expression are counted, the correlation of MEND counts to measured genes increases, while the correlation for MND decreases.

## Conclusion

Researchers want to know that their data is sufficient for the measurements they’re making. Here we show that, for the purpose of determining whether an RNA-Seq dataset is sufficient for accurately measuring expression of known genes, the fraction of relevant content of an RNA-Seq dataset (percent of MEND reads) varies substantially within and between cohorts. We confirm the relevance of MEND read counts to gene expression measurements by demonstrating that MEND read counts are correlated to the number of measured genes and total read counts are not.

This work was performed in pediatric samples as part of the development of our comparative RNA-Seq assay for pediatric cancer patients [6,12]. Since the factors that reduce the quality of RNA-Seq datasets (e.g. degradation, low sample volume, contamination, low base quality) are not specific to pediatric cancer samples, we predict that other kinds of RNA-Seq datasets would also show compositional variability. The MEND read counting tool is independent of species and genome version; it can be used on any bulk RNA-Seq dataset.

There are several reasons why a survey of this breadth has not been previously performed. Obtaining and processing clinical datasets from multiple sources is an intensive effort. Tumor datasets are usually controlled access, and obtaining the 48 cohorts we report on here required multiple legal agreements [25]. Analyzing read types requires genome-aligned reads; the files containing genome-aligned reads are large and are not generated when using the much faster pipelines that quantify gene expression via pseudoalignment. Large RNA-Seq cohorts such as GTEX and TCGA use consistent methods and exclude datasets that fail their stringent and consistent quality control [26,27]. They lack the kind of heterogeneity observed in our cohorts. In short, generating this data for more than 2000 datasets is time-consuming, expensive, and requires staff with a variety of expertises.

Measuring the number of MEND reads in a dataset is specific to the alignment parameters and gene model. We use Gencode v23, which is inclusive, defining more than 60,000 genes. By default, the aligner we use, STAR, defines reads that map to as many as 20 positions as mappable. If we changed our pipeline, asking STAR to exclude reads mapping to more than 2 positions and using a more conservative gene model with 30,000 genes, the same dataset would have fewer MEND reads due to the loss of reads that map too much or map only to regions newly defined as non-exonic.

People planning RNA-Seq experiments look for guidance on how much sequencing their experiment requires. For comparing gene expression measurements between datasets, ENCODE recommends a minimum of 30 million mapped reads [28]; the GEUVADIS consortium study had a minimum goal of 20 million reads [29]. However, of the 2078 datasets in this study with more than 30 million mapped reads, 16% contain fewer than 25% informative (MEND) reads. We speculate that these guidelines were not intended to include those datasets, some of which measure fewer than 100 genes.

Based on these results, we recommend that 1) publications reporting the results of an RNA-Seq study should report the number of MEND reads present in each dataset; 2) sensitivity studies should include read type fractions and report on the relationship between MEND reads and the measured outcome; and 3) sequencing depth recommendations should be based on MEND reads.

## Availability of supporting source code and requirements

Project name: MEND QC

Project home page: https://github.com/UCSC-Treehouse/mend_qc

Operating system(s): Platform independent

Programming language: Bash and R

Other requirements: Docker

License: MIT

RRID: if applicable, e.g. RRID: SCR_014986

## Acknowledgements

We acknowledge the work of all our colleagues at the Genomics Institute; the Computational Genomics Lab has provided an invaluable base for this work, allowing us to analyze large data sets relevant to pediatric cancer research. We thank Alejandro Sweet-Cordero and Alex G. Lee for valuable feedback on MEND analysis. We thank the many researchers who shared their sequence data [30]. Finally, we honor and thank all the children and adults who consented to donate their data to advance research in pediatric cancer.

## Supplemental figures

**Figure S1.**
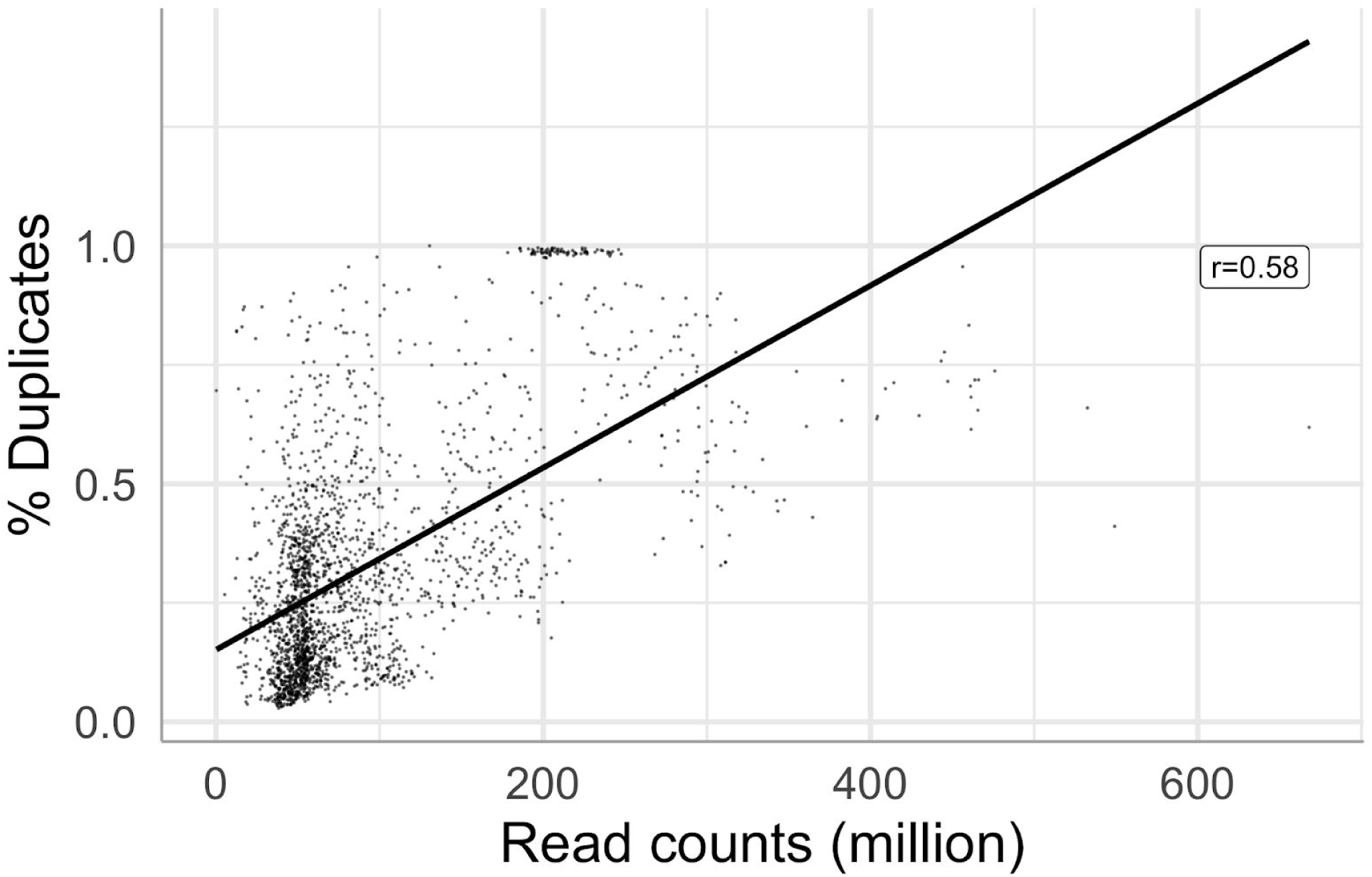
The percent of duplicate reads increases with the total number of reads in a dataset, but accounts for less than half of the variability. The Pearson correlation (r) between the values is 0.58 and explains 34% of the variability in the data (r^2^=0.336); n = 2179.

**Figure S2.**
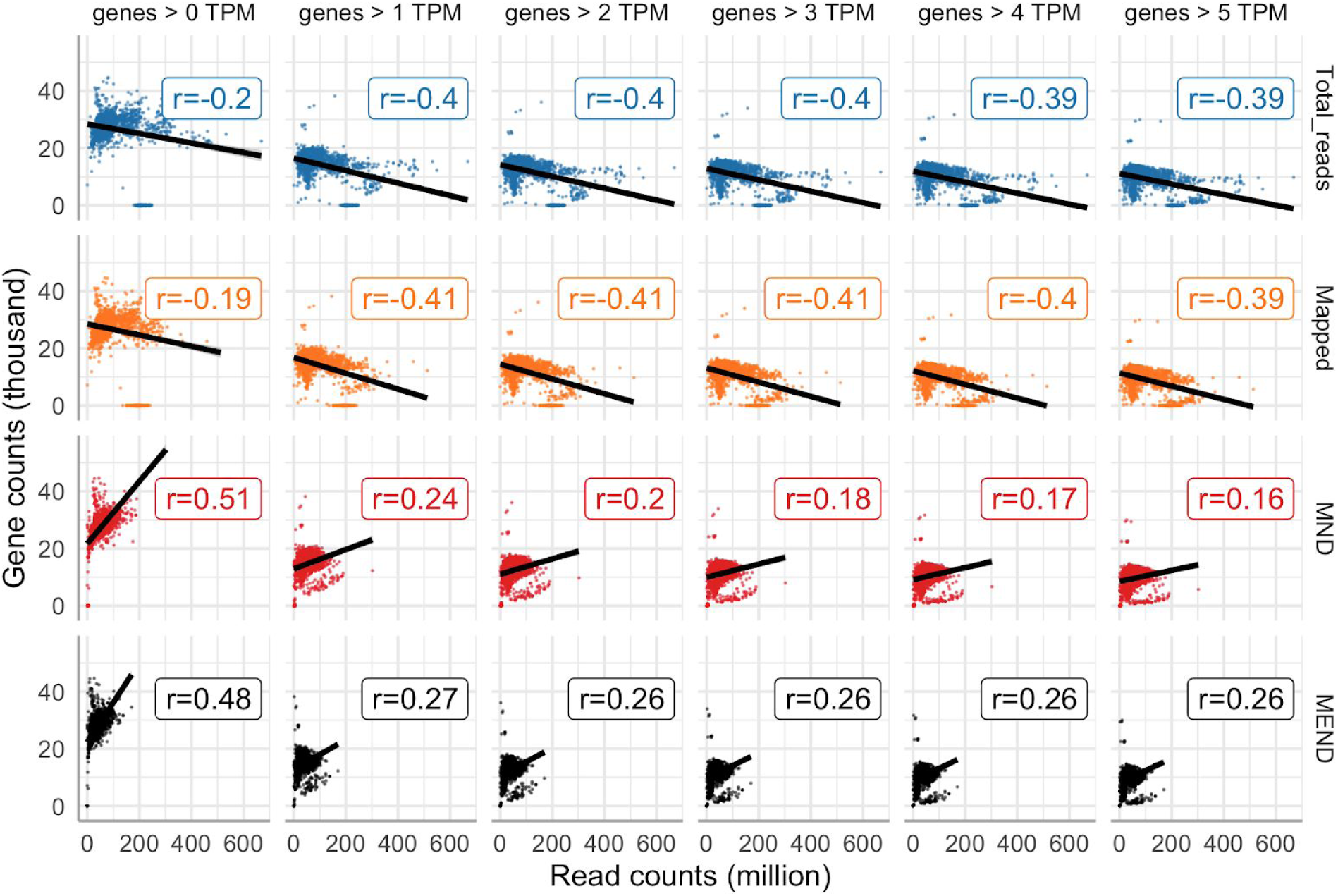
Of the four read types, MEND reads have the highest correlation to genes expressed above 1 TPM. The number of genes expressed (Y axis) above the threshold value (0, 1, 2, 3, 4, and 5 TPM, grouped by columns) are plotted against read counts (X axis). The type of reads counted (Total, Mapped, MND and MEND) are grouped by rows. The Pearson correlation (r) is shown for each combination of read type and gene threshold. All 2179 datasets are included in each plot.

Supplemental Table 1

https://docs.google.com/spreadsheets/d/1awKt5e3wYMWiliMwida1-8HLYrhUaOWP1GK26uV5ags/edit?usp=sharing

The accession numbers in Table S1 are the definitive sources of the RNA sequencing data. The DOI links to citations are provided for convenience. They may reflect the citation provided by the data provider, a citation we identified referring to the RNA sequencing data, or a citation we identified referring to the patient whose tumor was sequenced.

**Supplemental Table 2.**

| Name | Abbreviation | URL |
| --- | --- | --- |
| Short Read Archive | SRA | https://www.ncbi.nlm.nih.gov/sra |
| European Genome-phenome Archive | EGA | https://www.ebi.ac.uk/ega/home |
| St. Jude Cloud | SJC | https://www.stjude.cloud/ |
| Database of Genotypes and Phenotypes | dbGaP | https://www.ncbi.nlm.nih.gov/gap/ |
| Cavatica | Cavatica | https://cavatica.squarespace.com/ |

**Supplemental Table 3.**
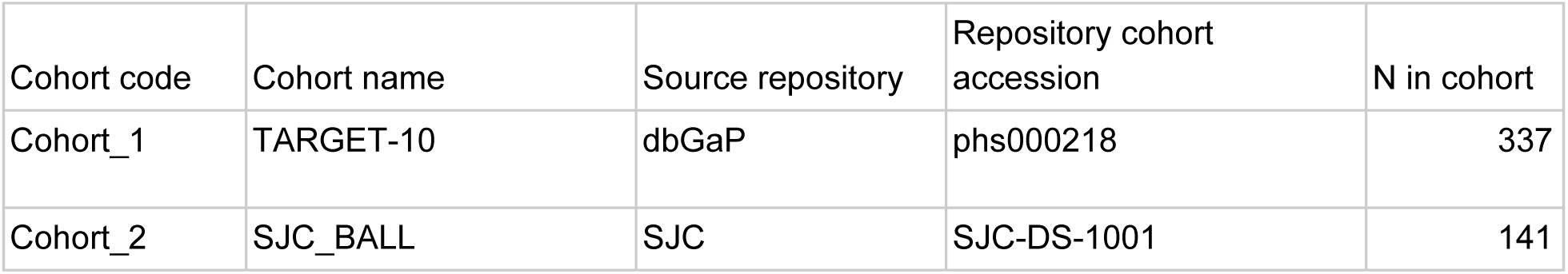
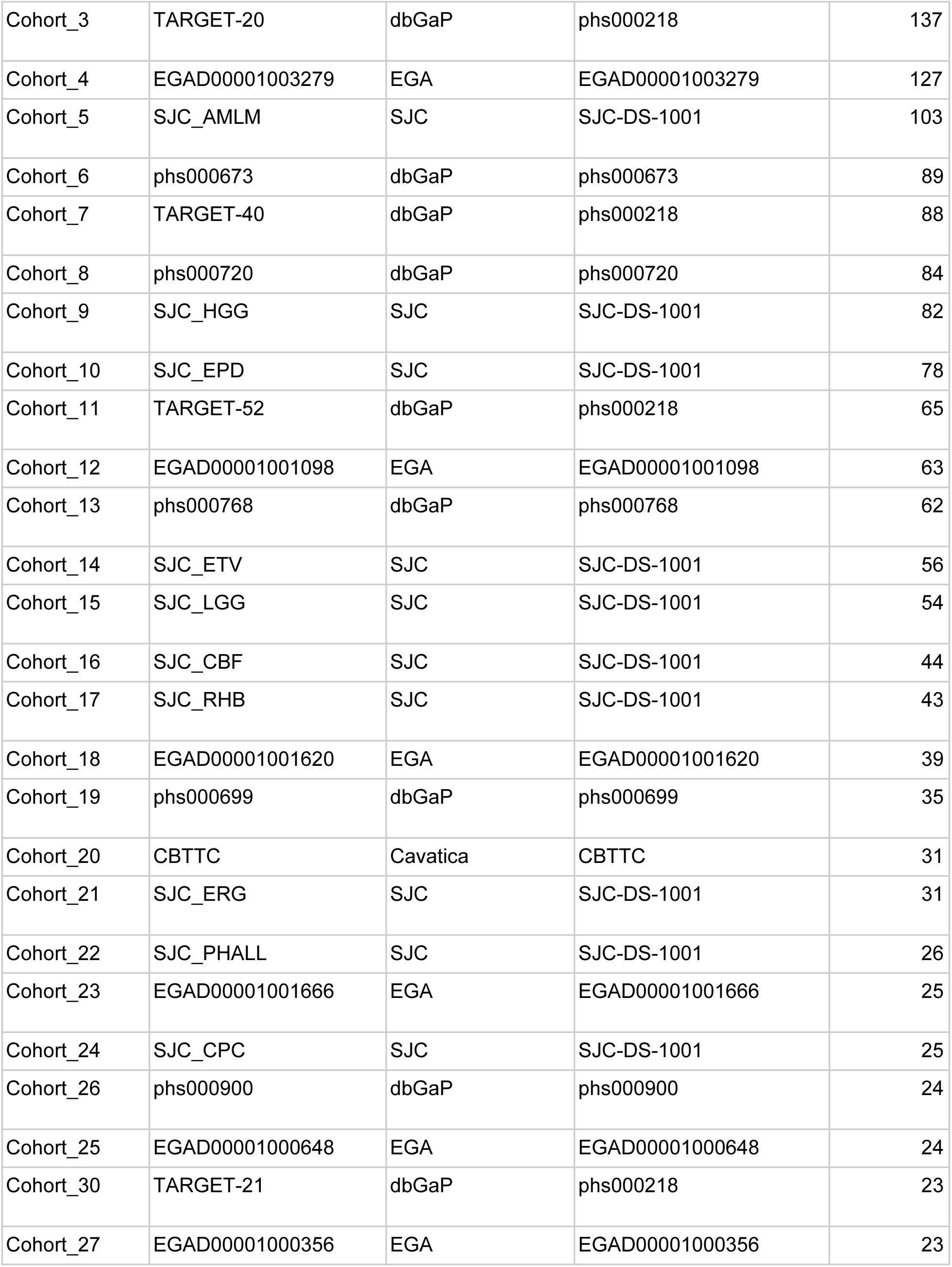
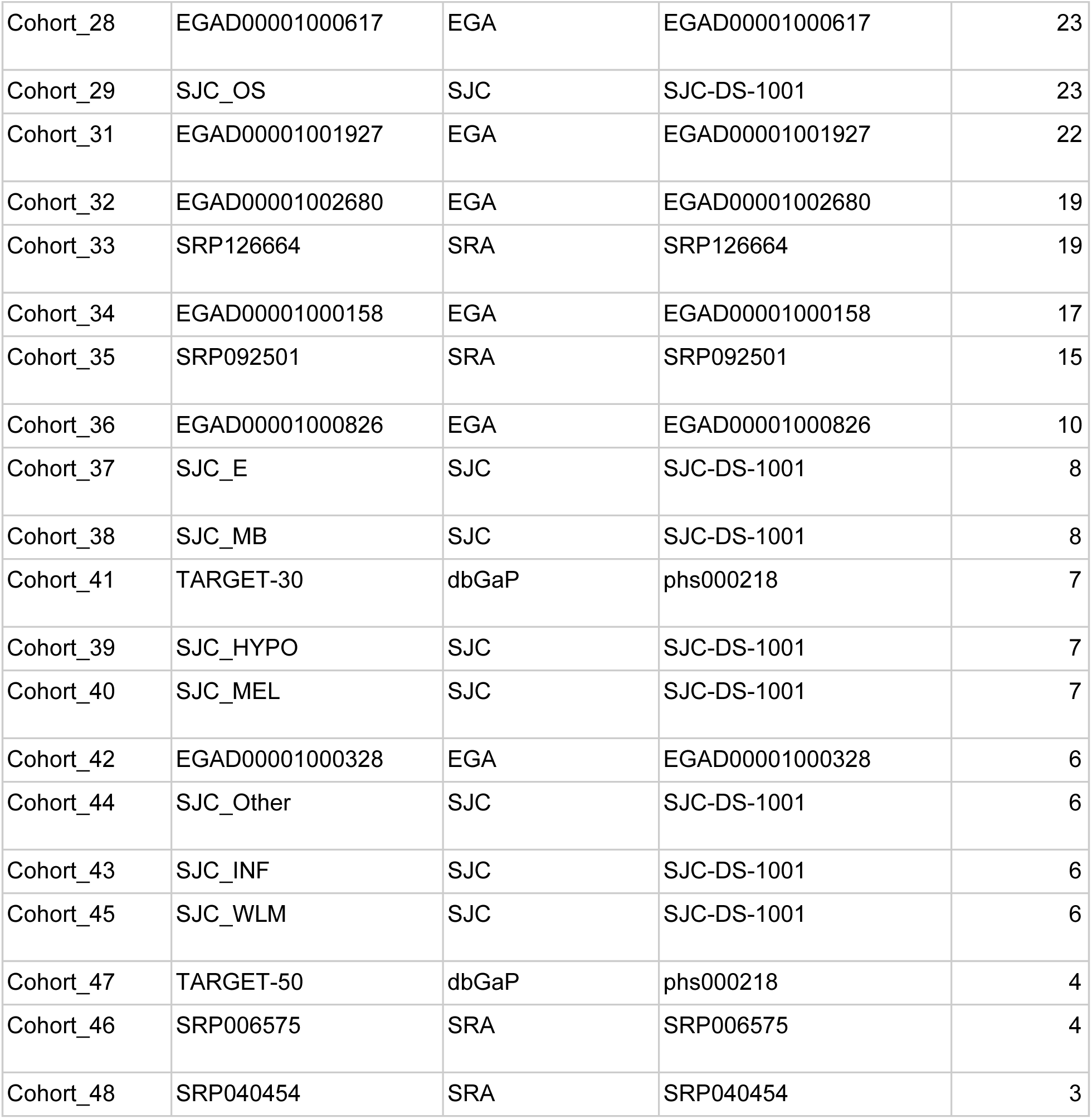

